# Establishing a DNA reference library for the identification of elasmobranchs in the Indian Ocean using Oxford Nanopore Sequencing

**DOI:** 10.1101/2025.06.03.657733

**Authors:** P.A.D.L. Anjani, Akshay Tanna, Emily Humble, W.M. Gayashani Sandamalika, Shaili Johri, Daniel Fernando

## Abstract

Elasmobranchs, comprising sharks and rays, are among the most threatened species globally. The first elasmobranchs were added to the Convention on International Trade in Endangered Species of Wild Fauna and Flora (CITES) in 2002. Since then, the number of species listed has increased to 151, highlighting the urgent conservation challenges faced by these species. In Sri Lanka, where over 100 species have been recorded, the demand for elasmobranch products drives significant export and import activities, complicating conservation efforts. Accurate species identification is crucial for fisheries monitoring, trade regulation, and conservation management. However, relying solely on morphological characteristics is challenging, especially since key identifying features are unavailable in certain derivatives like meat and oil in market samples. This study aims to develop a robust DNA reference library for elasmobranch species in Sri Lanka and to showcase the application of Oxford Nanopore Technology (ONT) as a reliable tool for fisheries monitoring and conservation. Using DNA barcoding and ONT sequencing, we analysed 49 elasmobranch species, focusing on the Cytochrome Oxidase I (COI) and 12S ribosomal ribonucleic acid (rRNA) genes to develop a DNA reference library. The creation of reliable barcode reference libraries is essential for effective species identification in biomonitoring because it facilitates the assessment of population trends and the effectiveness of fishing and trade regulations. We conducted a phylogenetic analysis to assess the robustness of reference sequences in species discrimination. All DNA reference sequences are publicly accessible through GenBank. This study enhances the capacity to monitor and regulate the elasmobranch trade in Sri Lanka and beyond.

## Introduction

Elasmobranchs (sharks and rays) are a group of cartilaginous fish that play a crucial ecological role in marine ecosystems [1]. These species typically exhibit low growth rates, late maturation, and long gestation periods [2,3,4]. Due to these biological characteristics, combined with increased fishing pressure, driven by global demand for their derivatives including shark and ray fins, gill plates, liver oil, and primarily domestic consumption of meat, elasmobranch populations are facing significant overexploitation and decline worldwide [5,6]. Currently, over one-third of sharks, rays, and chimaeras are threatened with extinction [7,8]. In response, Parties to the Convention on International Trade in Endangered Species of Wild Fauna and Flora (CITES) have taken measures to regulate the international trade of several elasmobranchs, most recently increasing the number of shark and ray species listed under CITES from 47 to 151 in 2022 [9]. These regulatory measures require tools to improve monitoring of elasmobranchs, from fisheries to trade, to enable better compliance and improve species management to prevent further decline.

Sri Lanka is an island nation with 23 operational major fish harbours [10]. The waters around Sri Lanka are home to over 100 elasmobranch species, of which 54 are shark species and 51 are ray species [12]. These species are both targeted and bycatch and utilised for local consumption and international trade [<u>10</u>; <u>13</u>]. The issue of illegal wildlife trade, particularly concerning shark fins and ray gill plates, has been a significant concern in Sri Lanka [14]. Between 2020 and 2021, customs authorities intercepted five illegal shipments involving over 300 shark fins and 800 <u>mobulid</u> ray gill plates, with some seizures surpassing 100 kg [15]. However, enforcement of fishing regulations remains challenging due to gaps in species-specific trade data and monitoring limitations at landing sites [15], particularly for derivatives. The mislabeling of shark and ray products further complicates monitoring, and Sri Lanka’s trade databases list only two commodity codes for these products, raising concerns about actual species in trade.

To address these challenges, Sri Lanka has developed National Plans of Action for the Conservation and Management of Sharks (NPOA-Sharks), following FAO guidelines to improve species identification, data collection, and trade monitoring; however, implementation has been limited. As a signatory to CITES, Sri Lanka is also required to conduct Non-Detriment Findings (NDFs) for the export of CITES-listed elasmobranch species, and as a signatory to CMS. However, the effective implementation of these measures is unclear. Improving species identification to expand data on species and derivatives in trade would be beneficial in supporting the implementation of CITES, CMS, NPOAs, or other conservation and management efforts. The strengthening and application of molecular and morphological data can enhance species verification, which alongside effective policy implementation, can aid enforcement agencies in better regulating elasmobranch fisheries and addressing illegal trade in Sri Lanka and globally. When considering molecular data, the most established method for species identification is DNA barcoding [17]. This technique typically involves DNA extraction, PCR amplification, Sanger sequencing, and sequence similarity analysis against a reference database [18]. One key limitation is that Sanger sequencing generates only a single sequence read, or electropherogram, for each sample’s PCR amplification product. When samples are contaminated or contain mixed-species templates, the electropherogram can become unreadable [19]. Since these DNA references obtained through this study will be used to identify elasmobranch products, such as oil and processed items, we proposed utilising a high-throughput sequencing (HTS) platform. These advanced technologies can generate amplicon consensus sequences with comparable or even higher accuracy than Sanger sequencing [20] and are more effective in distinguishing between contamination and mixed-species samples [21].

Oxford Nanopore technology (ONT] is one of the high-throughput sequencing methods, and its MinION sequencer is a small, portable, and affordable nanopore-based DNA sequencing platform. This technology offers several significant advantages over other high-throughput sequencing platforms, including the ability to produce long-read outputs, a low initial startup cost of around US$1,000, relatively simple and quick library preparation protocols, and the capability for rapid real-time analysis [<u>22</u>; <u>23</u>; <u>21</u>; <u>24</u>]. Given Sri Lanka’s need for reliable species identification services, its potential implementation in the government sector, and the high volume of import-export activities, we selected ONT for its cost-effectiveness, simplicity, and ease of use. In addition, this study focused on minimizing the cost by using the most cost-effective alternative equipment such as Bento Lab (Bento Bioworks, London, United Kingdom] instead of the Conventional PCR System and Rapid Sequencing Kit (Oxford Nanopore Technology, UK] instead ONT’s Ligation Sequencing kit and NanoDrop 1000 Spectrophotometer (Thermo Fisher Scientific, Waltham, USA) instead of Qubit fluorometer for quantification library. Therefore, this study produces DNA sequences from a cost-effective genetic pipeline which can be affordable for developing countries such as Sri Lanka. The findings of this study will aid in validating the visual identification of elasmobranch species. By establishing a DNA reference library using this data, we can create a reliable method for the preliminary identification of species at export points. This will ensure better implementation of CITES trade measures and allow for the disaggregation of trade data at the species level.

## Methodology

### Sample Collection

The tissue collection of Blue Resources Trust includes samples from over 100 chondrichthyan species. Tissue samples from 49 chondrichthyes species, landed at 28 locations across Sri Lanka, were selected over five years from 2018 to 2022 (Figure 1 and Table 1). To ensure representativeness, three tissue samples per species were selected for barcoding and molecular analysis. These samples were selected based on their IUCN Red List categories, the availability of adequate tissue samples, and their distribution across the country. Priority was given to species in threatened categories: Critically Endangered (CR), Endangered (EN), and Vulnerable (VU), as shown in Figure 2. A total of 95.9% of the sampled species belong to these threatened groups, while the remaining 4.1% fall under the Data Deficient (DD) and Near Threatened (NT) categories. Figure 3 presents the species composition by family for this study, with the highest number of species belonging to the family Dasyatidae. All these 15 families belong to 6 taxonomical orders under Elasmobranch.

**Fig 1.**
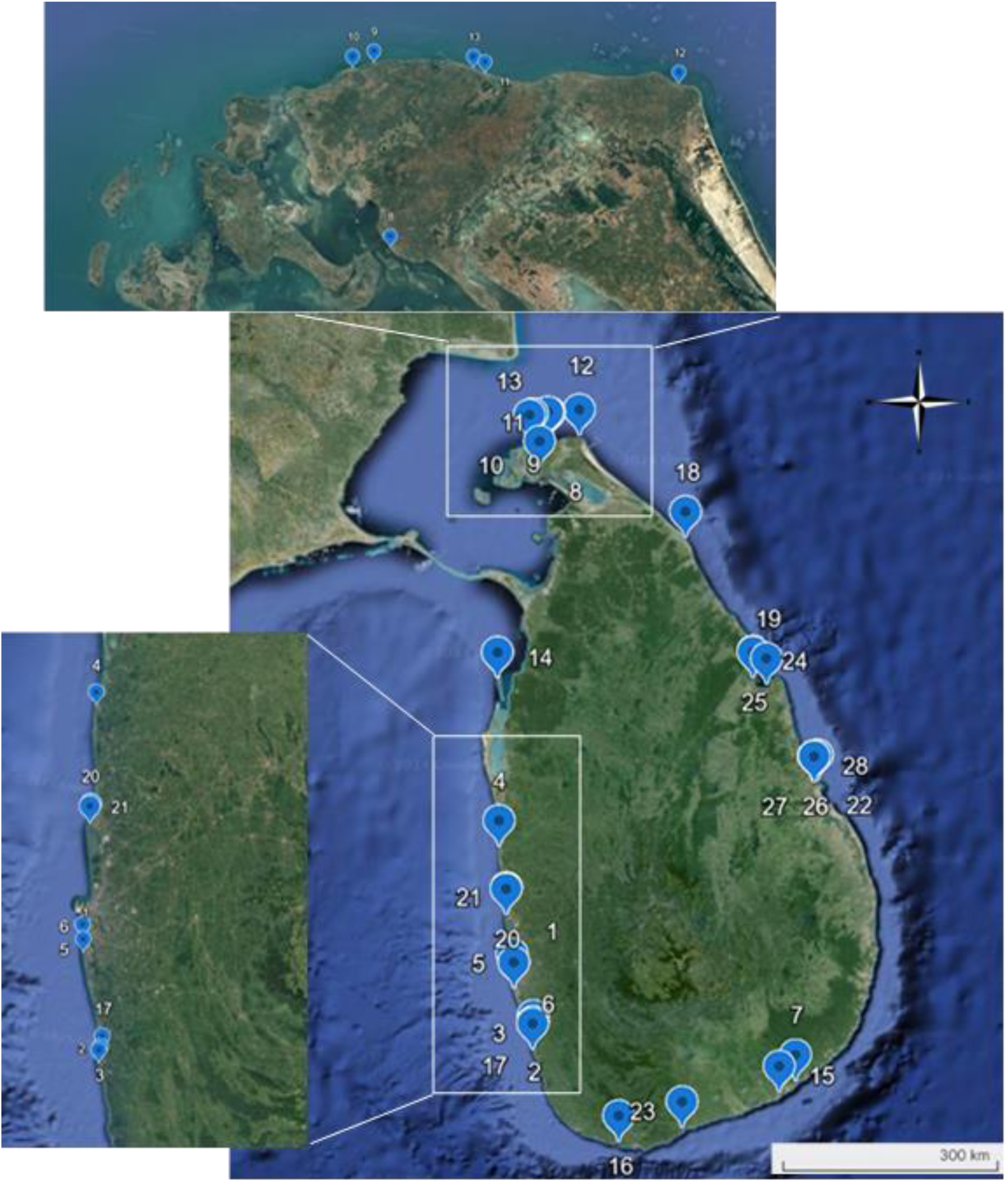
Elasmobranch sample collection sites in Sri Lanka. Samples were collected from 28 sampling sites round the Sri Lanka

**Fig 2.**
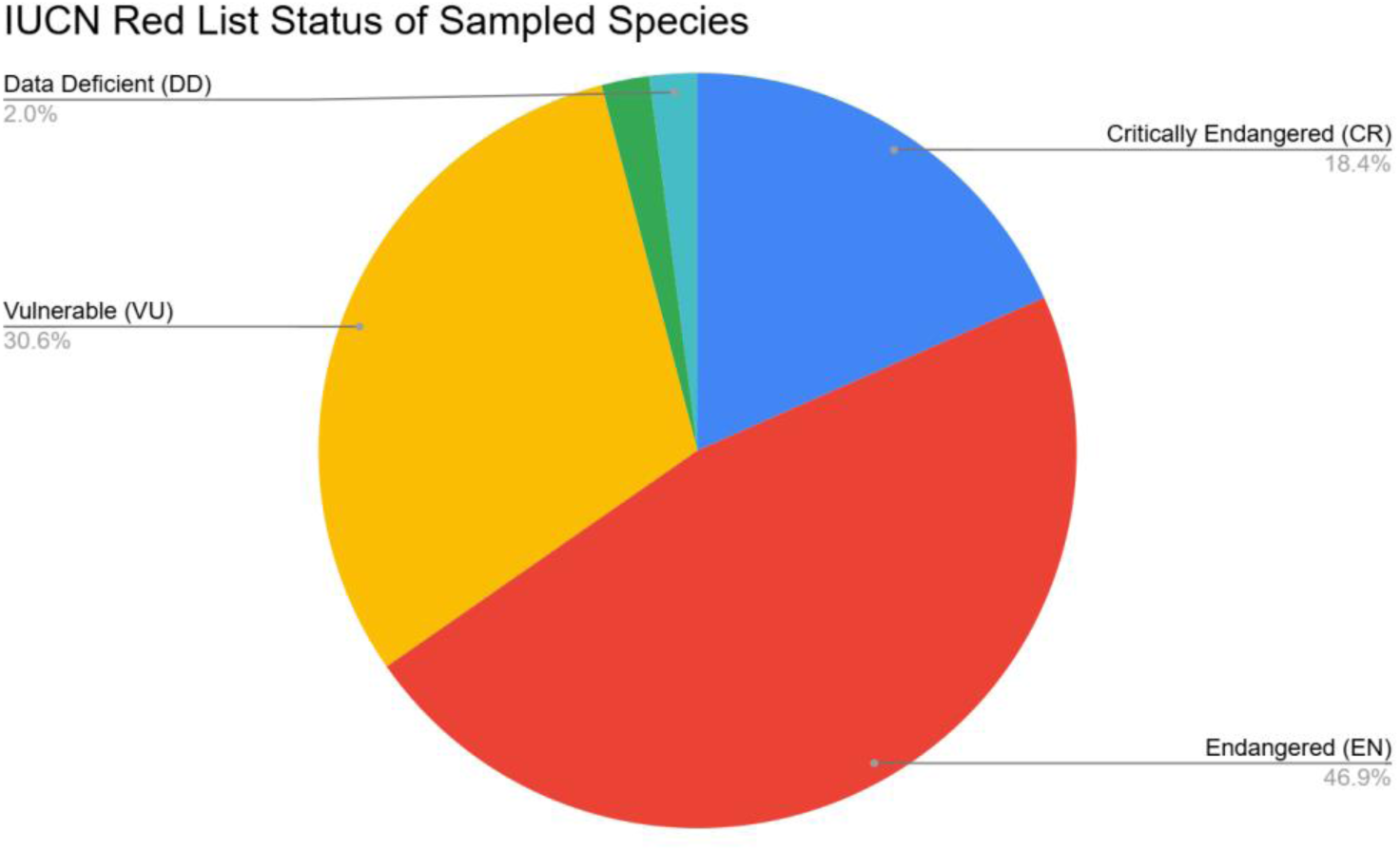
A pie chart showing the sampled species based on their IUCN status. All the sampling species belong to 5 IUCN categories; Critically endangered, Endangered, vulnerable, Data deficient and Least concern

**Fig 3.**
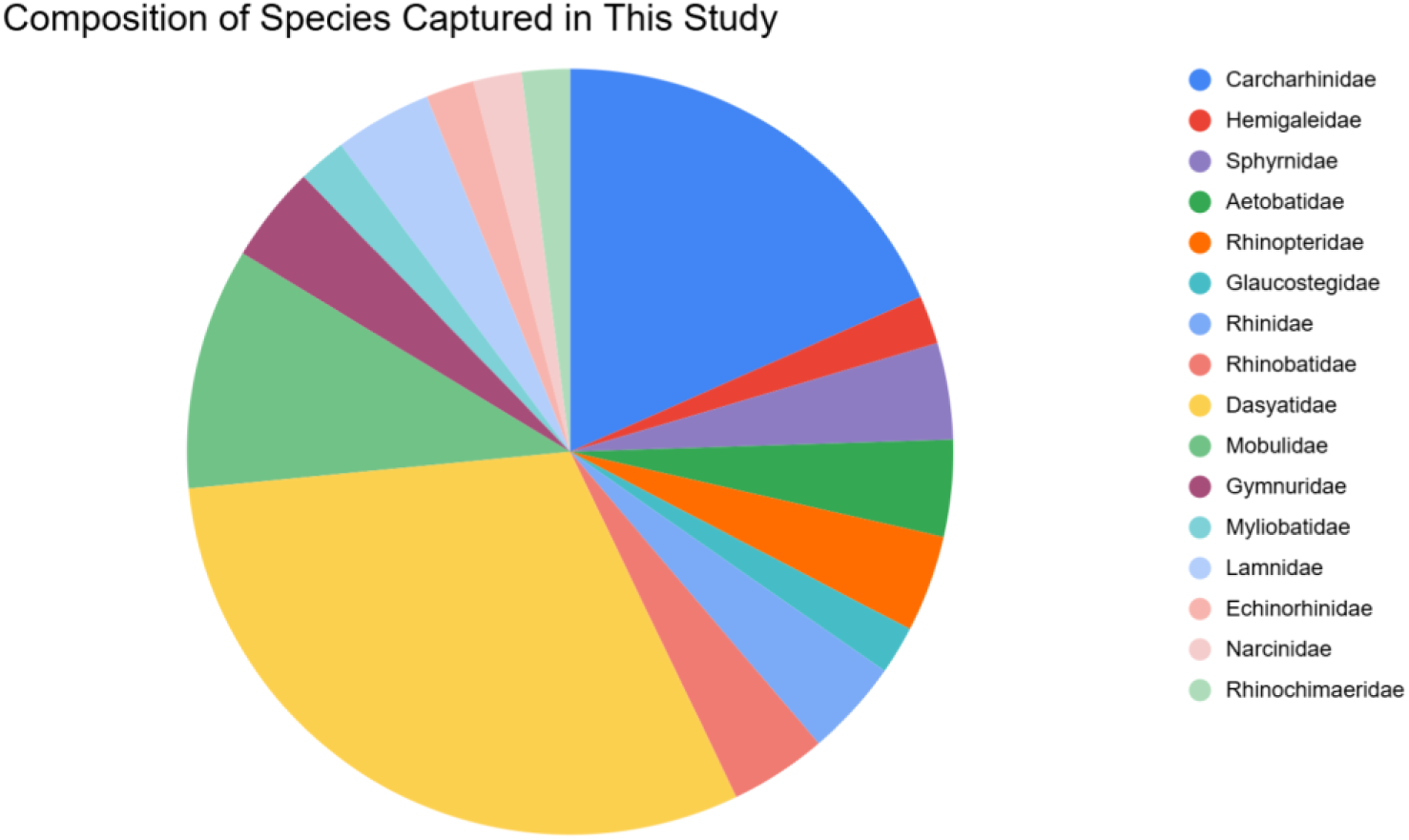
A pie chart illustrating the relative abundance of Families of different species used in this study as samples. All 49 species belong to 16 families according to their taxonomy

**Table 1.** List of species specimen numbers with major collection locations.

| **N**<br>**o** | **Site**<br>**Code** | **Reigon** | **Site**<br>**Location** | **Amount**<br>**of**<br>**samples** | **No** | **Site**<br>**Code** | **Reigon** | **Site**<br>**Location** | **Amount**<br>**of**<br>**samples** |
| --- | --- | --- | --- | --- | --- | --- | --- | --- | --- |
| **1** | ALS | West | 6.799773,<br>79.871102 | 3 | **15** | KFH | South | 6.21813,<br>81.336475 | 13 |
| **2** | BFH | West | 6.472438,<br>79.978596 | 2 | **16** | MFH | South | 5.948729,<br>80.449156 | 4 |
| **3** | BLS | West | 6.461017,<br>79.974955 | 1 | **17** | MGL | West | 6.505657,<br>79.978391 | 2 |
| **4** | CFM | West | 7.578421,<br>79.789994 | 13 | **18** | MLT | North | 9.268248,<br>80.821011 | 1 |
| **5** | CMB<br>D | West | 6.846209,<br>79.861898 | 2 | **19** | MUT | East | 8.463549,<br>81.266037 | 4 |
| **6** | CMB<br>L | West | 6.845756,7<br>9.861881 | 2 | **20** | NBO | West | 7.203319,<br>79.828919 | 9 |
| **7** | DLS | South | 6.279609,<br>81.426935 | 11 | **21** | NFM | West | 7.210335,<br>79.831248 | 1 |
| **8** | JFN | North | 9.654083,<br>80.013736 | 6 | **22** | PAP | East | 7.932189,<br>81.547725 | 10 |
| **9** | JKO | North | 9.805599,<br>79.975075 | 3 | **23** | TFH | South | 6.025065,<br>80.799172 | 1 |
| **10** | JMA | North | 9.798407,<br>79.957922 | 19 | **24** | TKB | East | 8.502291,8<br>1.192273 | 2 |
| **11** | JMY | North | 9.811282,<br>80.069923 | 1 | **25** | TKL | East | 8.498314,<br>81.191786 | 1 |
| **12** | JPP | North | 9.8282824,<br>80.234475<br>1 | 3 | **26** | VFH | East | 7.927459,<br>81.53019 | 20 |
| **13** | JUR | North | 9.814234,<br>80.059502 | 1 | **27** | VFM | East | 7.924404,<br>81.53001 | 2 |
| **14** | KBG | West | 8.498174,<br>79.781305 | 2 | **28** | VLS | East | 7.925731,<br>81.525729 | 8 |

Approximately 200 mg of muscle or fin tissue was collected from each species and preserved in absolute ethanol. Morphological species identification was carried out based on identification guides [<u>25; 26;</u> <u>27; 28</u>]. Each specimen was thoroughly documented by taking high-resolution photographs for reference and recording standard morphometric measurements, including total length (TL) and fork length (FL) for sharks and disc length (DL) and disc width (DW) for rays, and where possible, weight (W), using precise methodology and calibrated tools. Tissue samples were placed in 2 mL DNA LoBind Eppendorf® tubes filled with absolute ethanol. Samples were transported to the Genetics Laboratory at Blue Resources Trust, Sri Lanka and upon arrival, samples were stored at -20°C to maintain DNA integrity until further processing.

### Optimization of DNA extraction method

The extraction of genomic DNA from elasmobranch tissues requires methods that can effectively handle the unique biochemical composition of these organisms. For instance, the presence of inhibitory substances such as lipids and proteins can significantly affect the quality of extracted DNA, leading to challenges in downstream applications like PCR and sequencing [29] Various studies have highlighted the importance of optimizing extraction protocols to minimize contaminants and maximize yield [<u>30;31</u>;<u>32</u>] For example, the use of CTAB-based methods has been shown to effectively isolate DNA while reducing the presence of polysaccharides and phenolic compounds, which are common in various tissues [<u>33</u>;<u>34</u>]. This is particularly relevant for elasmobranch tissues, where similar contaminants may be present.

The efficiency of four different DNA extraction methods was evaluated: the Qiagen method using the DNeasy Blood and Tissue Kit (Qiagen, Valencia, CA, USA), the Macherey-Nagel NucleoSpin Tissue Kit (Macherey-Nagel, Germany), the HotShot DNA Extraction Kit (Bento Bioworks Ltd, London, United Kingdom), and the Chelex method (Bio-Rad, US). This evaluation included three types of tissue preservation methods: ethanol-preserved, dry, and frozen tissues. The full list of samples used for this is available in S1 Table.

DNA was extracted using a DNeasy Blood and Tissue Kit following manufacturer instructions (DNeasy Blood & Tissue Kits). For the HotShot method, the sample was cut into small pieces with the help of sterilized scissor and placed in tubes with 75 μl Alkaline Lysis Solution (25 mM NaOH, pH 12). Tubes were incubated at 95 °C for 30 minutes and cooled down to room temperature. After this cool-down, 75 μl of Neutralising Buffer (100 mM Tris-HCl, 0.5 mM EDTA, pH 8) was added. Extracted DNA was stored at -20°C. Chelex method was different from the manufacturer’s instructions. It was developed according to a previously described method [35], with some changes to improve optimisation. 200μl of 10% Chelex solution was added to the 1.5 ml Eppendorf tubes. These tubes were heated at 60°C for 10 minutes in a heat block, and samples were added to the hot Chelex solution. 15μl of proteinase K was added to the Chelex suspension with the sample and incubated at 55°C for 1 hour, vortexed, and again heated at 100°C for 15 minutes. The Eppendorf tubes were centrifuged at 10000 g for 10 seconds at room temperature. The supernatant was pipetted out gently to avoid extracting the Chelex resin and stored at -4°C. DNA was isolated from samples using the Macherey-Nagel NucleoSpin Tissue kit, according to manufacturer instructions and the extracted DNA was stored at -20°C until further use.

The study demonstrated that the amount of DNA extracted depended on both the extraction technique and the specific tissue type used. To further assess the success of each DNA extraction method, PCR amplification of the cytochrome oxidase subunit 1 (COI) gene was performed. The presence of clear and distinct PCR bands indicated successful DNA extraction and amplification. We compared the concentration and purity of isolated DNA, and the time and price of single extraction from elasmobranch tissues. By assessing the yield, purity, and integrity of the extracted DNA across different techniques, we found out the most suitable practices that can facilitate this research. All the statistics values obtained for determine the result listed in S2 Table As shown in Figure 4 and figure 5, the Qiagen DNeasy Blood & Tissue Kit and NucleoSpin Kit produced the most reliable results in terms of DNA concentration, purity, and PCR amplification success. However, these kits were the most expensive as shown in figure 6, costing US$ 215 and US$ 177 per kit, respectively, with a per-sample cost of US$ 4.30 for Qiagen kit and US$ 2.34 for NucleoSpin kit. As the result of this experiment, we found out column-based kits combined with ethanol-preserved tissues offer the most consistent and high-quality results for our research.

**Fig 4.**
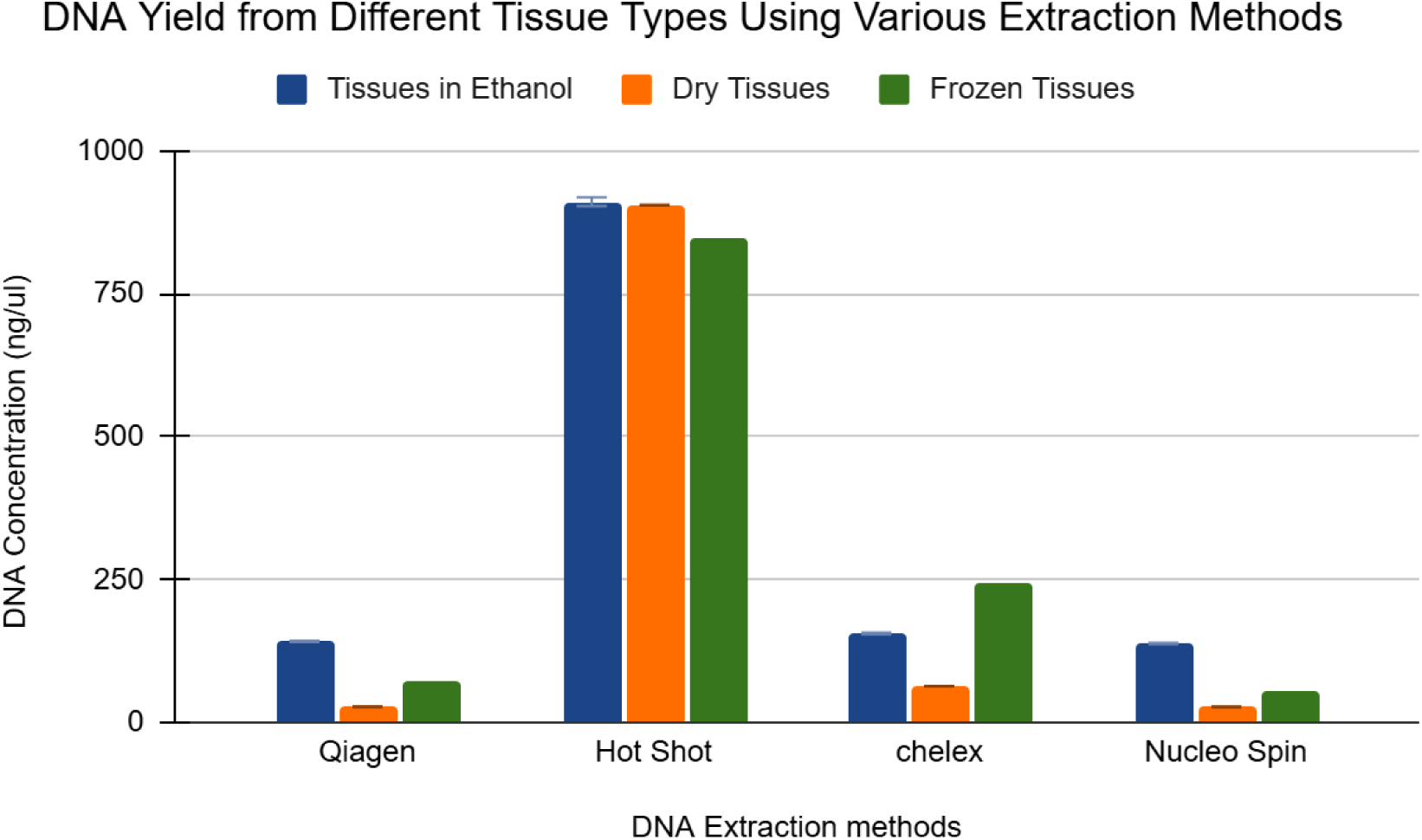
Concentrations of DNA extracted from different DNA extraction methods. Concentrations of DNA extracted from tissues in ethanol, dry tissues and frozen tissues using Qiagen method using DNeasy Blood and Tissue Kit (Qiagen), HotShot DNA extraction Kit (HotShot), Modified Chelex Method (Chelex) and Macherey-Nagel NucleoSpin Tissue Kit(Nucleo Spin). extraction methods

**Fig 5.**
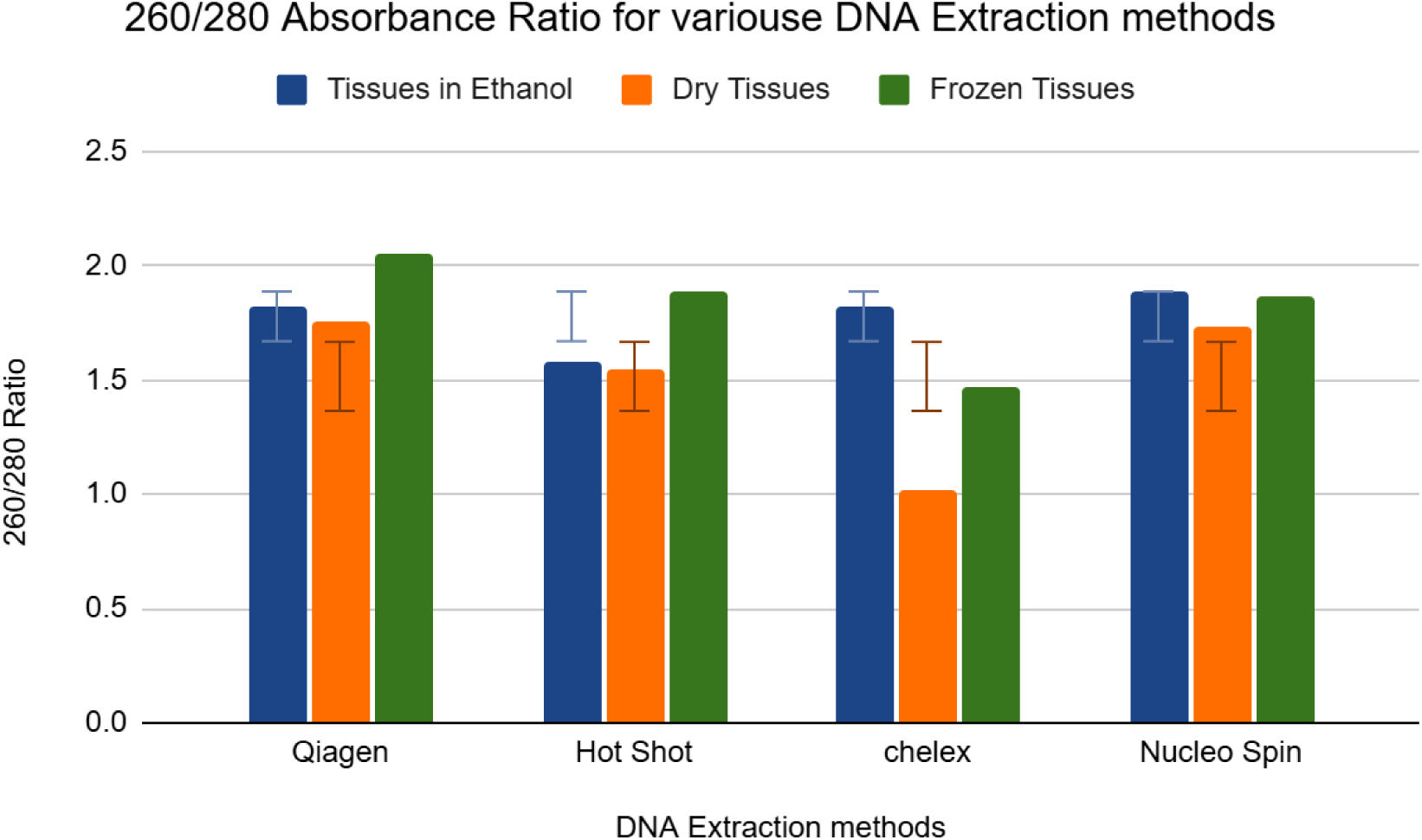
Purity ratio of DNA extracted from different types of samples. Purity ratio of DNA extracted from different types of tissues in ethanol, dry tissues and frozen tissues using Qiagen method using DNeasy Blood and Tissue Kit (Qiagen), HotShot DNA extraction Kit (HotShot), Modified Chelex Method(Chelex) and Macherey-Nagel NucleoSpin Tissue Kit (Nucleo Spin) extraction methods

**Fig 6.**
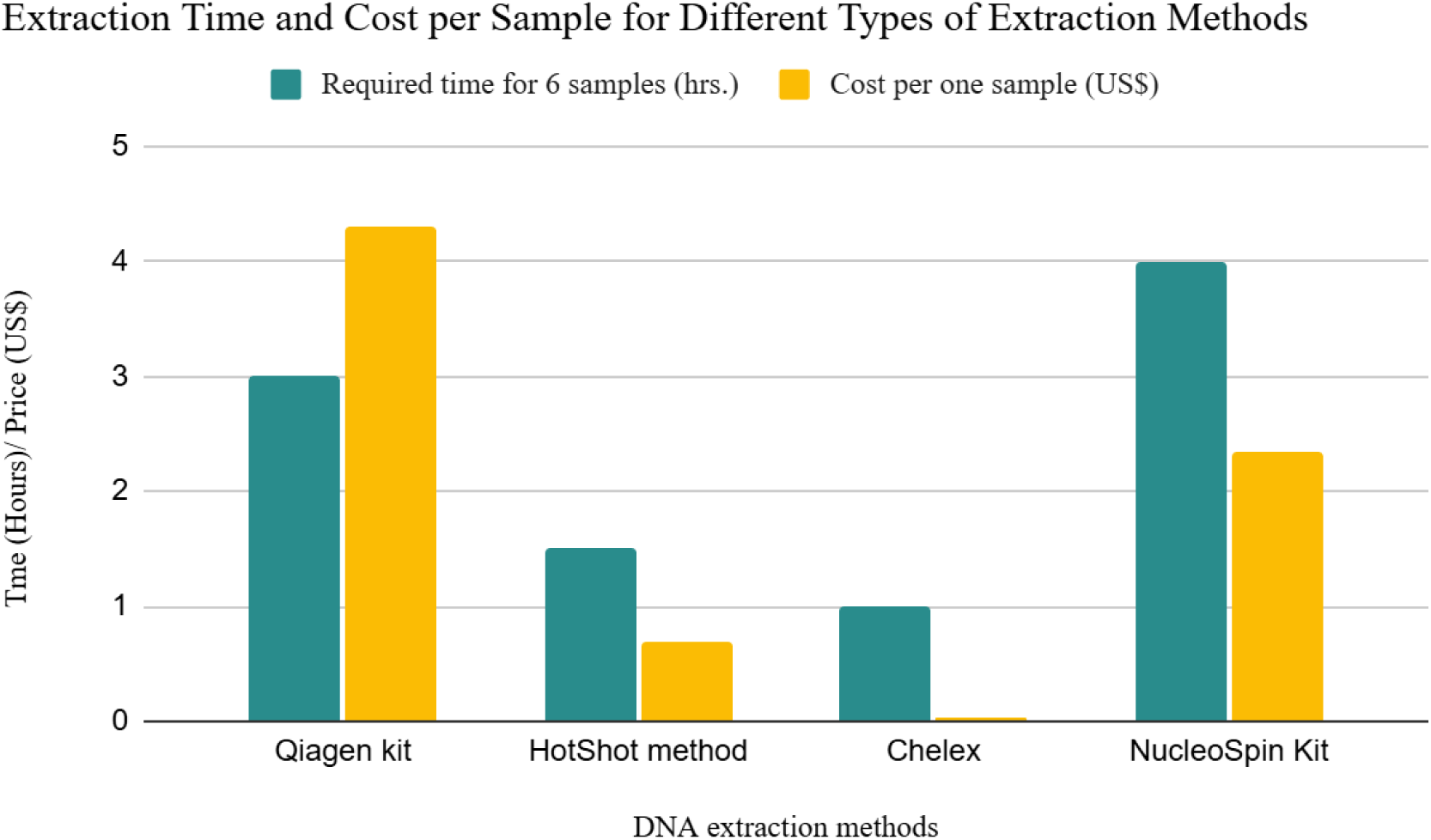
Extraction time and cost per sample across different DNA extraction methods. Bar plots show the time required to process six samples and the cost per sample (in USD) for each extraction method. Chelex and HotShot are the fastest and most cost-effective methods, while commercial kits such as Qiagen and NucleoSpin are more time- and cost-intensive

### DNA isolation

As we decided column-based kits combined with ethanol-preserved tissues give the best DNA for elasmobranch from above study, we isolated genomic DNA using the DNeasy blood and tissue kit (Qiagen, Valencia, CA), following the instructions of the manufacturer for isolate DNA to build up DNA reference library. A total of 20 mg of ethanol preserved tissue was measured from each sample, and excess alcohol was removed by blotting with tissue paper before the start of the lysis step. The DNA was eluted in 200 μl of elution buffer (AE buffer) at the final stage. Extracted DNA was quantified using a NanoDrop 1000 Spectrophotometer (Thermo Fisher Scientific, Waltham, USA) and the concentration and purity ratios were obtained. Extracted DNA was stored at -20℃ for further usage.

### PCR Amplification and visualisation

A 650 bp region of the cytochrome oxidase I (COI) gene from mitochondrial DNA was amplified using universal primers Fish F1 and Fish R1 primers [36] (Table 2). PCRs were carried out in a 20 μl reaction volume containing 1x FIREPol® Master Mix (Bento Bioworks, London, United Kingdom) and 100 ng of template DNA. The PCR profile described in Table 2 was repeated for 35 cycles, followed by a final extension for 10 min at 72 ℃.

**Table 2:**
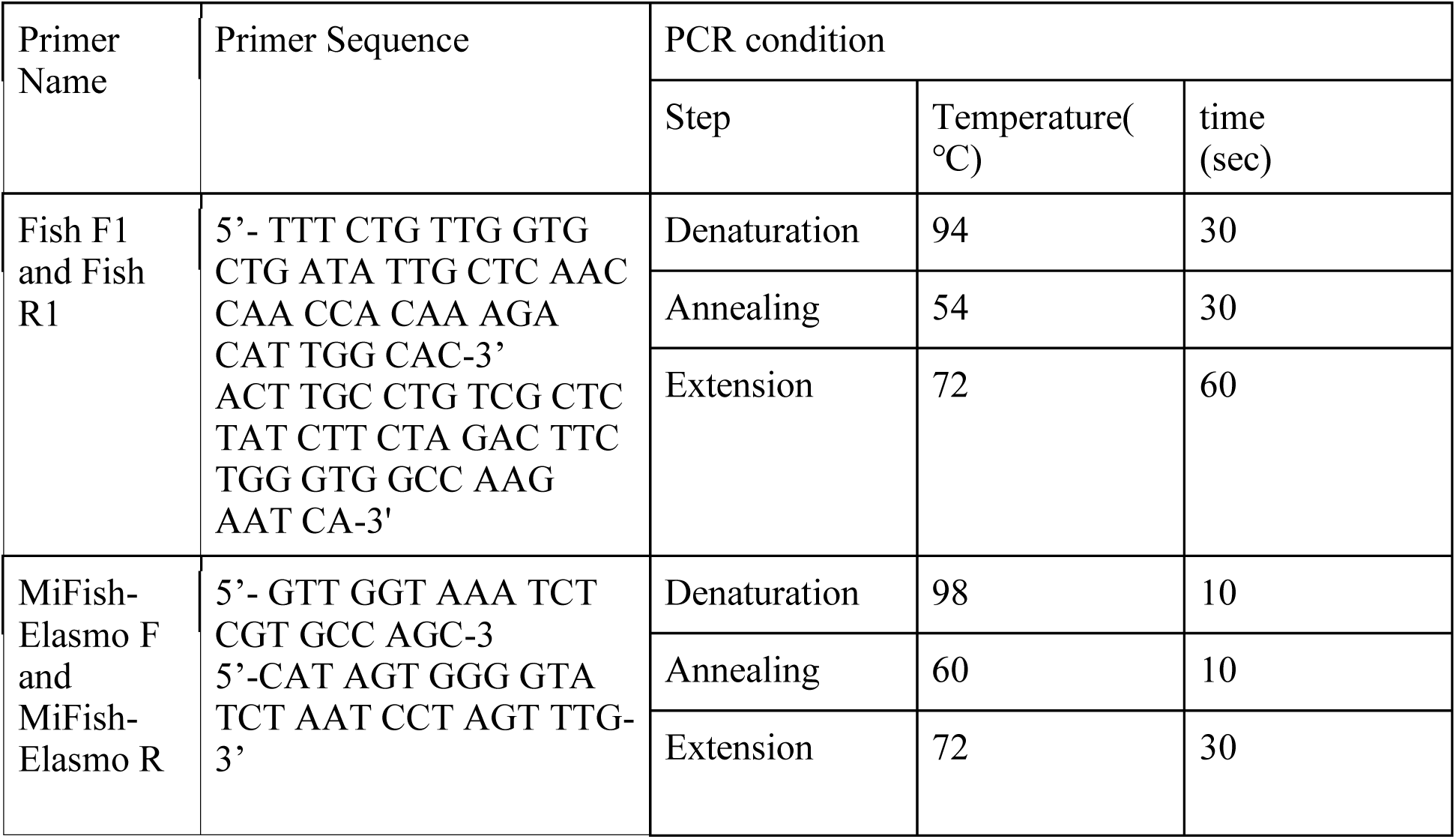
Sequences of primers used for this study and the time and temperature for each step.

Species that did not show amplification using Fish F1 and Fish R1 primers were subjected to amplification using MiFish-E primers, which are specific for elasmobranchs and amplify the 12S Ribosomal ribonucleic acid (rRNA) gene region [37]. The 12S rRNA region has been reported to be an excellent marker for the authentication of fish and seafood due to its mutation rate and acceptable length [38]. Approximately 200 bp of the 12S rRNA gene was amplified according to the conditions described in Table 2 while other gene amplified 655 bp amplicon. For both sets of primers, three PCR replicates were obtained from each sample to minimize stochastic variations. The three PCR products were then combined before proceeding with library preparation. We visualized PCR products on a 2% agarose gel with 0.5 x TBE buffer under blue light using the Bento Lab (Bento Bioworks, London, United Kingdom) with GelGreen® DNA stain (Bento Bioworks, London, United Kingdom).

### Library preparation and sequencing

The Rapid Barcoding kit (SQK-RBK110.24) from Oxford Nanopore Technologies, containing 24 unique barcodes, was used for the sequencing. PCR products were cleaned up using the HighPrep PCR magnetic bead solution (MagBio Genomics, Gaithersburg, USA), and DNA quantification was done using a NanoDrop 1000 Spectrophotometer (Thermo Fisher Scientific, Waltham, USA) for the cleaned-up products. Library preparation was done according to the manufacturer’s instructions for the Rapid Barcoding kit (SQK-RBK110.24) - Amplicon sequencing protocol by following PCR clean up, Amplicon DNA barcoding, Sample pooling and Adapter ligation. We used 15 samples for barcoding and pooling at once, considering the capacity of Flongle flow cells. Sequencing was carried out on a Flongle Flow Cell (R10.4.1) using the MinION Mk1B, where 200 - 400 fmol of the library was loaded onto the flow cell via the sample port. Ten libraries were sequenced for approximately 20 hours to obtain at least 30,000 raw reads per run.

### Data Analysis

#### Consensus Building

GitHub workflow (https://github.com/anjani95l/BRT_ElasmoGenetic_2022_anjani.git) was used for data analysis. We basecalled raw sequence data using Guppy v6.5.7 (ONT) with the base-calling model “dna_r9.4.1_450bps_hac.cfg”. Quality control was carried out using MinIONQC according to the previously described method [39] protocol, followed by demultiplexing with Guppy. Reads with q scores less than 7 and read lengths that did not equal the amplicon length (with a 50 bp buffer) were filtered using Nanofilt. Read clustering, consensus forming, and consensus polishing were carried out using NGSpeciesID. Reads were first clustered according to similarity using ONclust. Consensus sequences were generated for each cluster using SIMD Partial Order Alignment (SPOA), which also conducts error corrections, resulting in more accurate consensus sequences. Further polishing of the consensus sequence was done using Racon (<u>21</u>). Primers were also removed at this step. Species identities were assigned to each consensus sequence using a nucleotide BLAST search v2.15.0 [40]. Species assignment was based on multiple criteria, including percent identity, query coverage, E-values from BLAST results, and phylogenetic tree placement, to ensure accurate identification. Identity percentages from blast results were recorded because they give the highest percentage identity for a set of aligned segments to the same subject sequence (Library Guides: NCBI Bioinformatics Resources: An Introduction: BLAST: Compare & identify sequences).

#### Phylogenetic tree reconstruction

To validate species identification and assess genetic relationships, phylogenetic trees were constructed using the Neighbour-Joining (NJ) method in MEGA11 software v11.0.13. The phylogenetic analyses were performed separately based on the genetic markers used for sequencing, as gene selection plays a crucial role in tree topology. Sequences obtained from the cytochrome c oxidase subunit I (COI) and 12S rRNA genes were analyzed separately, leading to the formation of two primary groups. To ensure accurate species categorization, the dataset was further divided based on taxonomic classification and sample size, resulting in the construction of five distinct phylogenetic trees. These included (1) species from the orders Carcharhiniformes and Lamniformes, (2) species from the family Dasyatidae within the order Myliobatiformes, (3) remaining species within Myliobatiformes, (4) species from the orders Rhinopristiformes and Torpediniformes, and (5) a phylogenetic tree exclusively for species identified through the 12S rRNA gene. This approach allowed for a more refined resolution of species relationships within each taxonomic group and ensured the reliability of our genetic pipeline for species identification. For the phylogenetic tree construction, MEGA11 software v 11.0.13 was used, and the alignment of sequences was done using the Clustal W algorithm.

Then, the phylogenetic trees were constructed using the Neighbour-Joining (NJ) method, and the reliability of the Neighbour-Joining method was tested by bootstrapping with 1000 replications of the data [41]. The maximum composite likelihood model was used for NJ analysis, with a transition: transversion ratio. Reference sequences were selected based on the results obtained from the NCBI BLAST search. Species with the highest query coverage and percent identity were chosen as reference specimens for the phylogenetic tree construction.

Morphological validation was conducted using voucher specimens, identified based on key external characteristics as mentioned in the methodology. Any discrepancies between genetic and morphological identifications were re-evaluated, with misidentified specimens corrected based on both genetic placement in the phylogenetic tree and expert morphological examination. Samples that showed unexpected clustering in the phylogenetic trees were examined closely to eliminate the possibility of sequencing errors, hybridization, or the presence of cryptic species. All validated DNA sequences were submitted to GenBank, enhancing a publicly accessible reference library for elasmobranch species in Sri Lanka.

## Results

We were able to identify sequences from 46 species out of 49 species, while three species presented challenges in amplification or identification. *Neoharriotta* pinnata did not amplify with either the Fish F1R1 primers or MiFish-E primers, likely due to primer mismatches. *Rhinobatos annandalei* was successfully amplified using the 12S rRNA gene. However, the absence of reference sequences for this gene in GenBank prevented direct species confirmation. *Carcharhinus amblyrhynchoides* produced inconclusive results, as it did not align with the correct species in NCBI BLAST and exhibited unexpected placement in the phylogenetic tree, suggesting possible misidentification or incomplete reference data.

### Tissue sample collection intensity

A total of 7,141 tissue samples were collected over 10 years (2012–2022) from 45 landing sites and markets across Sri Lanka. Figure 7 illustrates the distribution of these samples around the country, highlighting the variation in sample collection by region. The highest number of samples, 3,519, was collected from the East Coast, while the South Coast contributed the fewest, with only 428 samples. The species with the largest sample size was *Neotrygon indica*, accounting for 1,107 samples out of 7,141. In this study, our samples were selected based on the criteria outlined in the sample collection section. Approximately 2% of the total samples were obtained from each region, allowing for a balanced representation of genetic diversity and species distribution across the study area. This approach helped minimize regional bias and provided a more comprehensive understanding of the species’ genetic characteristics across different geographic locations.

**Fig 7:**
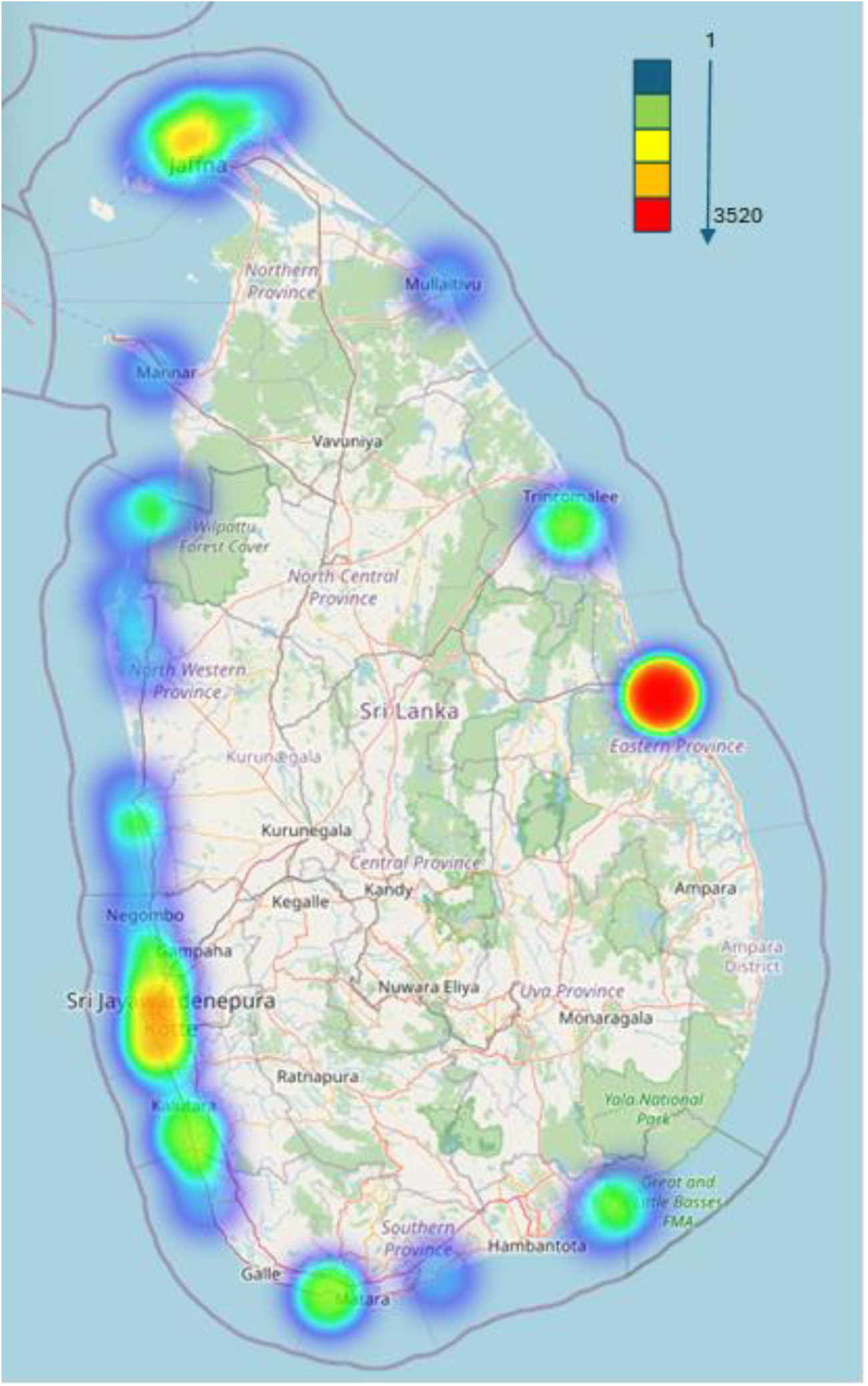
Distribution of collectuin of elasmobranch tissue samples from 2012 to 2022. The heat map provides a clear representation of sample collection intensity across different regions, with colors indicating varying levels of species count. Blue represents areas with a low species count, Green indicates moderate species count, Yellow shows higher species count, Orange highlights regions with high species count, and Red represents areas with a very high species count.

### Phylogenetic Analysis

The findings presented in Figure 8A reveal that eight species from the order Carcharhiniformes form a distinct clade, separate from the two species of the order Lamniformes. This separation is reflected in the genetic distances shown in the phylogenetic tree, where species from Carcharhiniformes cluster closely together, indicating a higher genetic similarity, while the Lamniformes species are positioned further apart, suggesting a greater genetic divergence. *Isurus oxyrinchus* and *Isurus paucus* belong to Lamniformes, commonly referred to as Mako sharks, display significant morphological similarities. The primary features used for differentiation include variations in fin structure and body coloration; however, these characteristics may not be readily apparent in some instances, such as processed carcasses, complicating species identification. This study proves, DNA barcoding provides a reliable method for distinguishing between these two species. In Figure 8B, a phylogenetic tree is presented, illustrating the relationships among 15 sampled species within the family Dasyatidae. The species *H. leoparda, H. uarnak, and H. undulata*, previously classified as part of the *Himantura urnak* species complex due to their close morphological similarities, are grouped together in this analysis. Notably, the 45K sample of *H. undulata* is positioned within a separate clade, indicating a significant divergence from its reference species. This sample exhibits a difference of 60 base pairs from the reference, with no gaps in the alignment, suggesting its distinctiveness. The *Pastinachus* species, including samples *P. bleekeri* (85Y, 87Y, 86Y), demonstrate a high degree of similarity to the reference species *P. uarnacoides*, with 675 out of 683 base pairs being identical and only an 8 base pair difference in the alignment. Additionally, the phylogenetic tree depicted in Figure 8C incorporates sampled species from the families Aetobatidae, Rhinopteridae, Mobulidae, and Gymnuridae. It is particularly challenging to identify species within the Mobulidae family when they are processed into pieces for market sale. However, the phylogenetic tree reveals that all five species exhibit distinct lineages, confirming that they can be differentiated through DNA barcoding. In Figure 8D, the sequences from *R. australiae* and *N. timlei* are derived from the COI gene, while the remaining species sequences are derived from the 12S rRNA gene. Figure 8E consists of species from different families and orders, with a phylogenetic tree constructed based on the 12S rRNA gene sequence of these species. *E. bruscus*, the samples 73U and 75U cluster closely, supporting their correct species assignment. They are positioned alongside *E. cookei* (LC146082.1), confirming only their placement within the Echinorhinidae family. *C. leucas* (142R.1, 144R, 143R.1) and *C. limbatus* (94B) align separately from their respective GenBank references in this phylogenetic tree.

**Fig 8.**
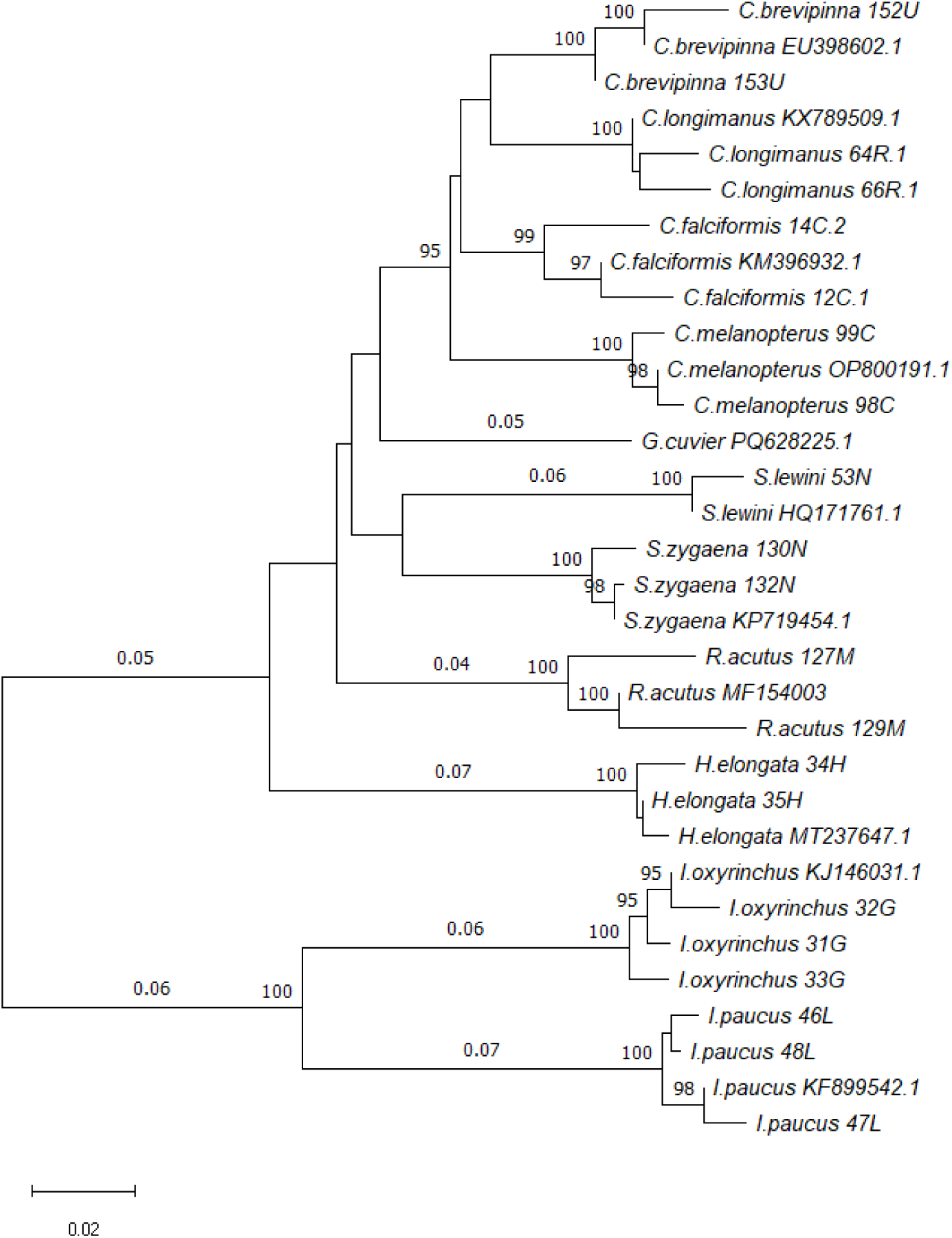
Maximum Likelihood (ML) phylogenetic tree of species belongs to order Carcharhiniforms and Lamniformes with *Galeocerdo cuvier* as an outgroup. This is based on partial nucleotide sequences of the cytochrome oxidase 1 (COI) mitochondrial gene sequences. The sample sequences are indicated by species name_sample number_english letter while reference sequences are indicated by species name_Acession number within the phylogenetic tree. Branch lengths representing genetic distances and bootstrap values indicating the statistical confidence of each node. Nodes with high bootstrap values (≥95) suggest strong phylogenetic relationships.

**Fig 8B.**
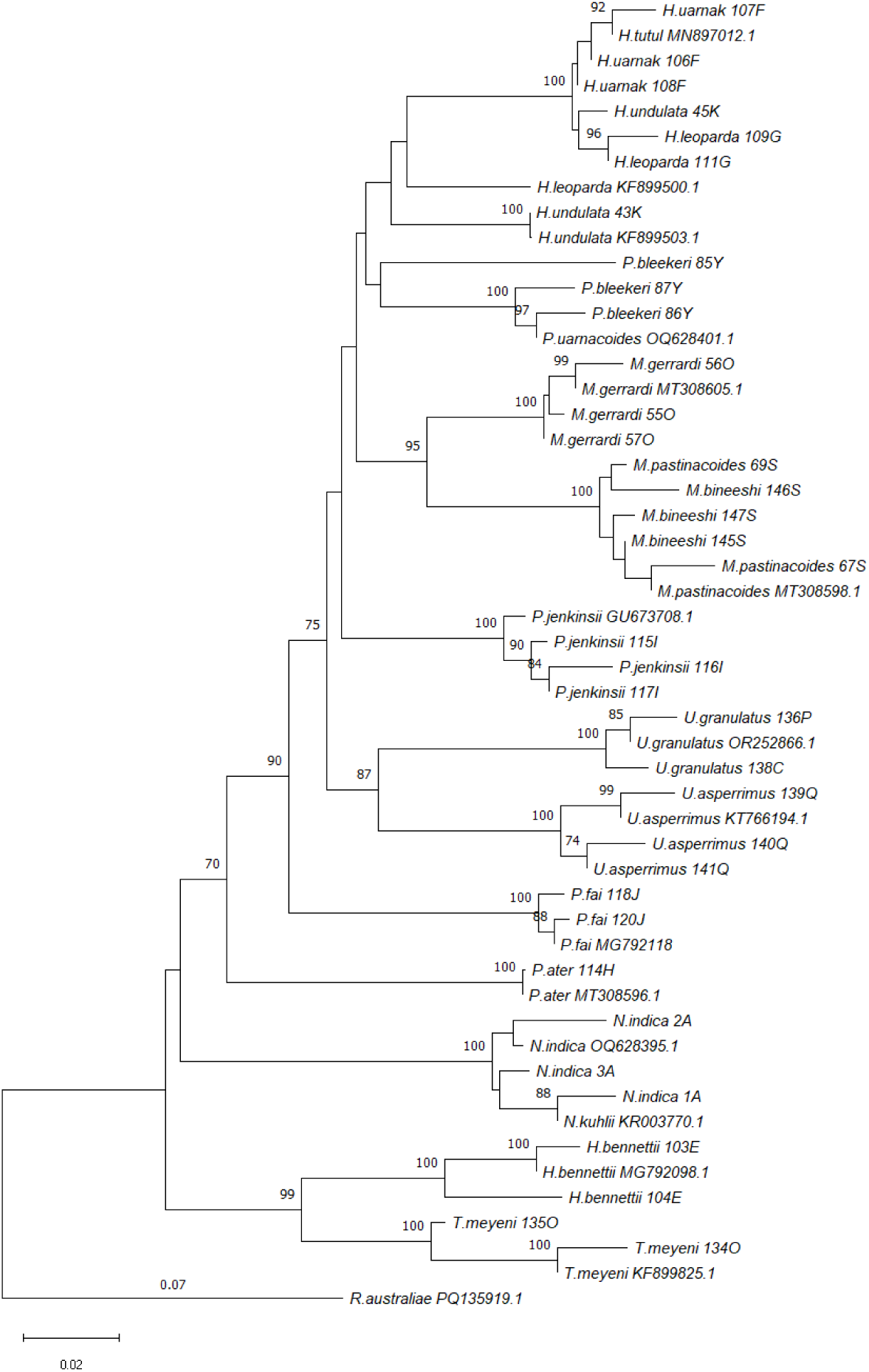
Maximum Likelihood (ML) phylogenetic tree of species belongs to family Dayasatidae of Order Myliobatiformes and *Rhinobatus australiae* as an outgroup. This is based on partial nucleotide sequences of the cytochrome oxidase subunit 1 (COI) mitochondrial gene sequences. The sample sequences are indicated by species name_sample number_english letter while reference sequences are indicated by species name_Acession number within the phylogenetic tree. Branch lengths representing genetic distances and bootstrap values indicating the statistical confidence of each node. Nodes with high bootstrap values (≥95) suggest strong phylogenetic relationships.

**Fig 8C.**
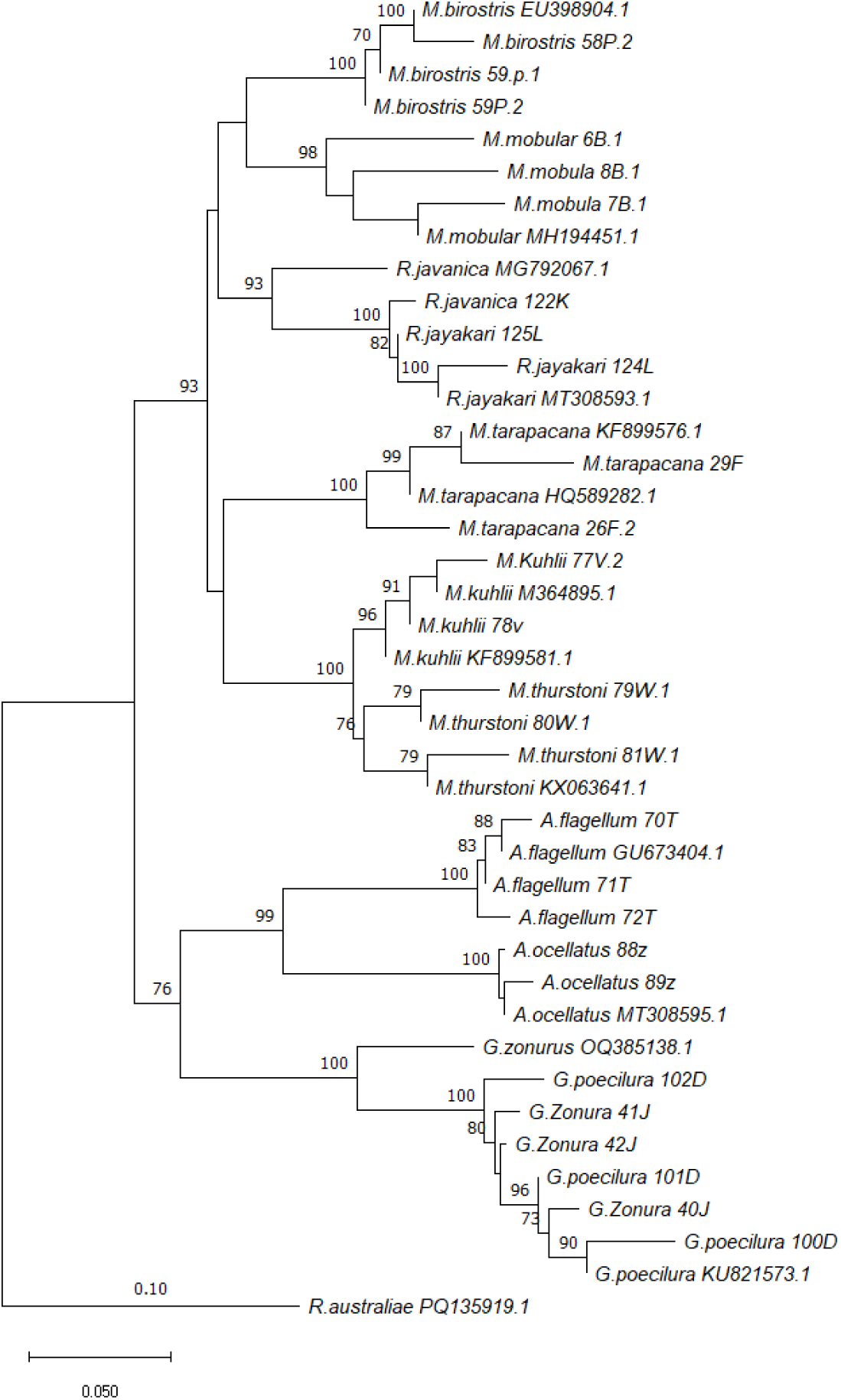
Maximum Likelihood (ML) phylogenetic tree of species belongs to Order Myliobatiformes and *Rhinobatus australiae* as an outgroup. This is based on partial nucleotide sequences of the cytochrome oxidase subunit 1 (COI) mitochondrial gene sequences. The sample sequences are indicated by species name_sample number_english letter while reference sequences are indicated by species name_Acession number within the phylogenetic tree. Branch lengths representing genetic distances and bootstrap values indicating the statistical confidence of each node. Nodes with high bootstrap values (≥95) suggest strong phylogenetic relationships.

**Fig 8D.**
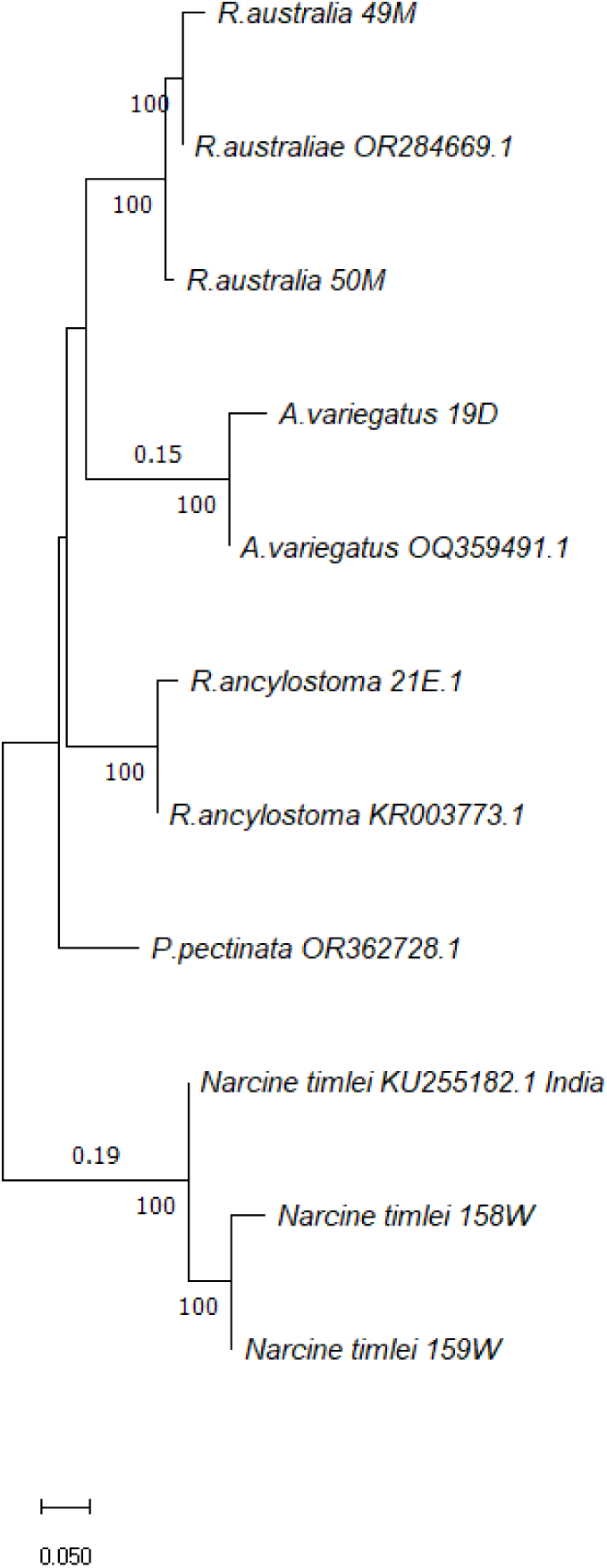
Maximum Likelihood (ML) phylogenetic tree of species belongs to Order Rhinopristiformes and Torpediniformes. This is using 12S Ribosomal ribonucleic acid (rRNA) and partial nucleotide sequences of the cytochrome oxidase subunit 1 (COI) mitochondrial gene. The sample sequences are indicated by species name_sample number_english letter while reference sequences are indicated by species name_Acession number within the phylogenetic tree. Branch lengths representing genetic distances and bootstrap values indicating the statistical confidence of each node. Nodes with high bootstrap values (≥95) suggest strong phylogenetic relationships.

**Fig 8E.**
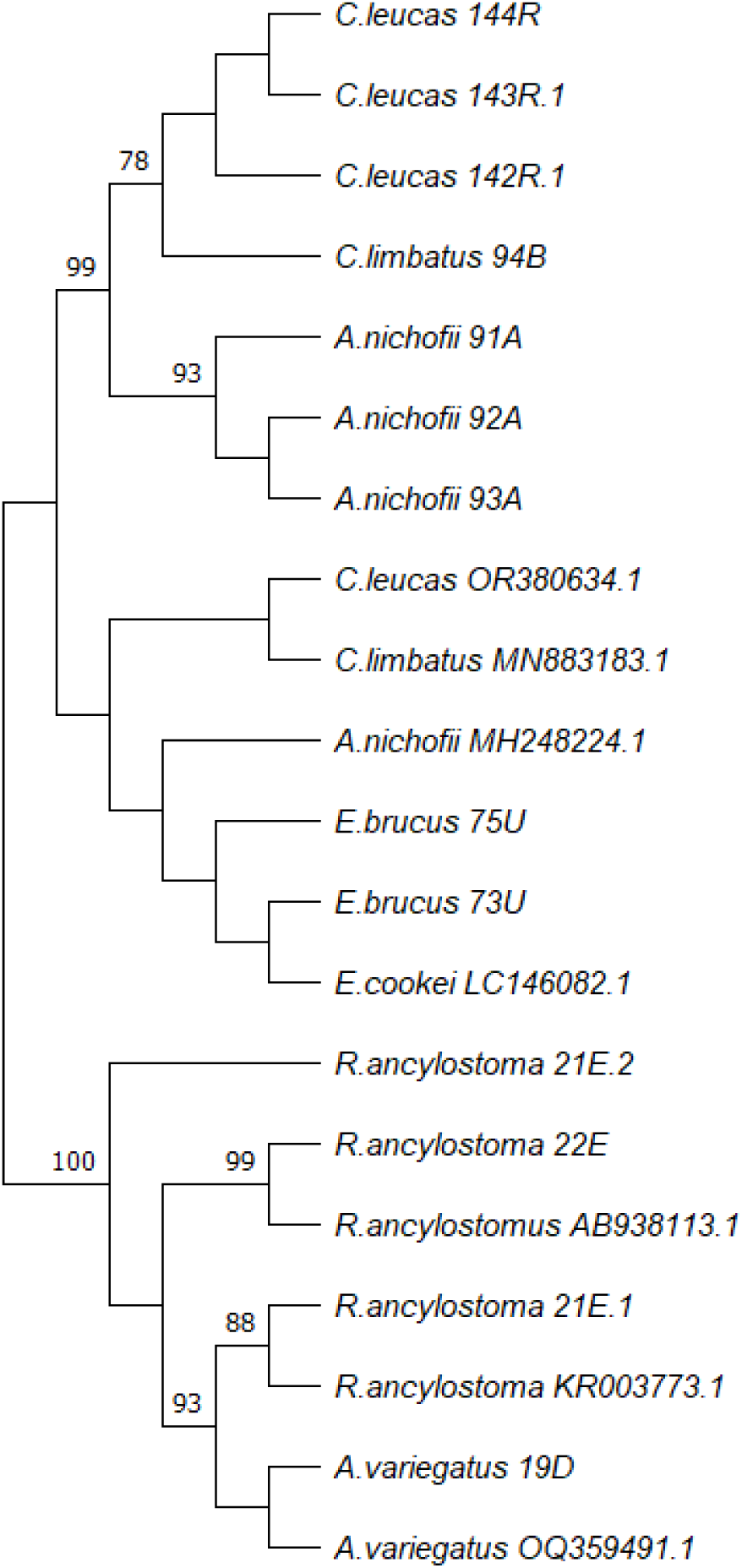
Maximum Likelihood (ML) phylogenetic tree of species amplified their 12S rRNA gene for the study. This tree has species who do not amplified well from Fish primers, and they belong to different Elasmobranch families. The sample sequences are indicated by species name_sample number_english letter while reference sequences are indicated by species name_Acession number within the phylogenetic tree. Branch lengths representing genetic distances and bootstrap values indicating the statistical confidence of each node. Nodes with high bootstrap values (≥95) suggest strong phylogenetic relationships, while lower confidence values at certain nodes suggest areas where additional sequencing or taxonomic revision may be needed.

## Discussion

In this study, we employed DNA metabarcoding techniques to analyze a total of 48 elasmobranch species inhabiting the Indian Ocean around Sri Lanka. By sequencing the Cytochrome Oxidase I (COI) and 12S rRNA genes, we developed a comprehensive reference library, which is vital for future taxonomic and ecological studies. Our findings revealed that we were able to identify 46 species using the COI gene and an additional 8 species through the 12S rRNA gene.

The significance of this study lies in its contribution to validation of elasmobranch species diversity in Sri Lanka through genetic tools. This effort therefore complements prior studies [42], and aligns with other genetic studies [<u>10;43</u>], which also aimed to catalogue the elasmobranch species in this region. This study has established the basis for a genetic databank of Sri Lankan elasmobranchs. Moreover, the establishment of a DNA reference library for elasmobranchs, as initiated in this study, represents a crucial step towards enhancing national research capacity. Such efforts could help overcome issues around shark conservation due to resource management challenges [44]This study also aligns with the establishment of other elasmobranch DNA reference libraries, such as ELASMO-ATL [45] and ELASMOMED [46], which have been demonstrated to be invaluable for elasmobranch conservation efforts across the globe.

The introduction of Oxford Nanopore technology for the identification of elasmobranchs represents a significant advancement in molecular ecology in Sri Lanka, which in turn can support monitoring of products in trade to implement management. This third-generation sequencing method allows for the concurrent processing of multiple samples, enabling the analysis of up to 24 specimens in a single run of this study. The increasing accuracy and performance of Oxford Nanopore sequencing, especially its ability to produce highly accurate consensus barcodes relative to other next-generation sequencing methods, highlights its potential utility in biodiversity assessments and species monitoring [47].

In this context, rapid and real-time sequencing techniques, such as those offered by Oxford Nanopore Technology, could serve as crucial tools for the identification of shark and ray products, facilitating timely interventions in export monitoring. Moreover, the portability of the equipment utilized in this study enhances its applicability in field studies and onboard identification, allowing researchers and enforcement agencies to conduct assessments in situ. This combination of technological advancement and practical application has the potential to significantly strengthen efforts in the conservation of vulnerable Elasmobranch species. However, it is important to note that despite the promising potential of ONT, significant challenges exist in its practical implementation. As it stands, acquiring ONT reagents and equipment is incredibly difficult for countries with restrictive import regulations, such as Sri Lanka, combined with the logistical barriers practised by Oxford Nanopore Technologies with respect to shipping destinations, cold chain requirements, and expirations of products. This limitation significantly hampers the ability to integrate such advanced sequencing techniques into routine monitoring and enforcement activities, particularly in countries where these species are vulnerable to illegal trade. Until these logistical challenges are addressed, the full potential of ONT in conservation efforts may remain underutilized. Despite these obstacles, the combination of technological advancement and practical application could, if accessible, greatly enhance efforts to monitor vulnerable elasmobranch species.

The study of DNA optimization in this research demonstrates that the choice of DNA extraction method and tissue preservation type significantly influences the quality and quantity of extracted DNA for molecular applications. The results showed that the Qiagen DNeasy Blood & Tissue Kit and NucleoSpin Kit were the most effective methods for obtaining PCR-amplifiable DNA from ethanol-preserved, dry, and frozen tissues. In contrast, the HotSHOT method, while yielding high DNA concentrations, failed to produce PCR bands, indicating poor DNA quality. The Chelex method produced amplifiable DNA only from ethanol-preserved tissues, demonstrating limited applicability across preservation types. Dry tissues yield the lowest quality and quantity for every extraction method. This is primarily due to the drying processes, particularly air-drying or heat-drying, which can lead to the fragmentation of DNA strands caused by enzymatic activity before the tissue fully dehydrates [31]. Once DNA is fragmented, recovering high-molecular-weight DNA becomes more challenging, impacting downstream applications such as PCR and sequencing. Drying can also cause cross-linking between DNA and proteins, especially in tissues with a high protein content [48]. These complexes are more difficult to break apart during extraction, resulting in lower yields and purity.

The highest DNA quantity came from the HotShot method, which utilizes an alkaline lysis buffer to efficiently release DNA. Alkaline extraction produces DNA yields equivalent to or higher than the spin-column method because it contains NaOH, which rapidly breaks down cell membranes and nuclear envelopes [49]. However, DNA degradation can also occur due to the same alkaline solution, as alkaline conditions can cause DNA strand breaks and base modifications, especially if the heating step is prolonged or not precisely controlled [49].

This results in fragmented or partially degraded DNA, which shows low DNA quality. Most importantly, HotSHOT and Chelex methods do not remove proteins, salts, or inhibitors due to a lack of purification systems, unlike other column-based methods [29]. These contaminants can interfere with downstream applications like PCR, even if the DNA quantity is high.

Under the optimization of DNA extraction method in this study shows ethanol preserved tissues are the best to use for DNA extraction because of various characteristics in ethanol such as it kills decomposing microorganisms and it removes water from the tissues, slowing down enzymatic processes and it denatures both the DNA and DNA-degrading enzymes preventing further enzymatic degradation [<u>50</u>;<u>51</u>]. Column-based DNA extraction methods are considered among the best due to their efficiency, purity of output, and adaptability for different sample types. This happened due to silica membranes in columns, which selectively bind DNA, allowing impurities to be washed away efficiently. Considering all these factors we used Qiagen DNeasy blood and tissue kit and Ethanol preserved elasmobranch for DNA extraction for DNA barcoding study.

The findings of this DNA barcoding study highlight the complexities associated with species identification among elasmobranchs, particularly when relying on molecular techniques. While the COI gene is widely accepted as a robust marker for DNA barcoding, some elasmobranch species did not yield successful amplifications. Consequently, the 12S rRNA gene was employed for identification. However, we encountered a significant challenge due to the limited availability of 12S rRNA reference sequences within the NCBI database, which adversely affected the clustering accuracy in our phylogenetic analyses, as shown in Figure 8E. This lack of reference data reduced the accuracy of our phylogenetic analyses, as the unavailability of comprehensive and current sequences for certain species and genes limited our ability to make accurate genetic comparisons. Several different primer sets can be used to amplify loci such as 12S, and because there are several different metabarcode regions within it, it is difficult to know if a sequence annotated as “12S” contains the marker fragment of interest [52]

Previous studies suggest that correct identification of species is expected when sequence similarity is above 97.5% for the COI gene [53]. Specimens used in this study were evaluated for sequence similarity for reference species and all species had over 97.5% of sequence similarity except the *Echinorhinus bruscus* specimens. As shown in Figure 8E, the specimens identified morphologically as *E. bruscus* clustered more closely with *E. cookei* and there is minimal divergence between the *E. bruscus* samples to the reference species. Previous study also suggest that *Echinorhinus* species found in Sri Lanka may represent an undescribed species [43]. Neighbour-joining analysis shown in Figure 8B showed the relationship between *Himantura urnak* and *Himatura tutul*. These two species showed a close relationship within one clade. The colour pattern of H. tutul described by Borsa [<u>54</u>;<u>55</u>] is the same as that of our specimens collected as H. uarnak. Previouse study also reported that the NADH2 gene shows H. uarnak from Sri Lanka is closely related to H. tutul from Indonesia, a finding that is validated by our results for the COI gene analysis [43]. In the same phylogenetic tree, *Neotrygon indica* and *Neotrygon kuhlii* are shown to be closely related, clustering together in a single clade. This close relationship is due to the species previously referred to as *Dasyatis kuhlii* (Müller & Henle) in Sri Lanka [56, 57, 58], which is now classified under the genus *Neotrygon* (Last & White 2008). In Figure 8B, *Maculabatis bineeshi* is closely related to *Maculabatis pastinacoides*, and *Pateobatis bleekeri* is closely related to *Pateobatis uarnacoides.* This clustering occurs due to the unavailability of reference sequences for *Maculabatis bineeshi* and *Pateobatis bleekeri* in GenBank. As a result, samples from these two species formed a phylogenetic tree with the most closely related species within their respective families. Additionally, the absence of reference data in GenBank for *M. bineeshi* may be attributed to the fact that the species was only described in 2016 (Manjaji-Matsumoto and Last 2016). Previous studies also highlighted the lack of data related to the 12S rRNA gene on batoid fish [61]. However, the methods used to obtain, filter and archive sequences obtained from public databases are often poorly documented [52], while the scope for updating and reusing these data is limited. Improvement in these aspects will increase the reliability, flexibility, and transparency of metabarcoding protocols.

Despite these limitations, the high recovery rate (94%) of species sequences demonstrates the robustness of our methodological approach. Future studies may overcome these gaps by optimizing primer design, increasing sequencing depth, or incorporating additional genetic markers to improve detection sensitivity for these species.

These findings underscore the importance of thorough taxonomic investigations and the expansion of genetic databases to support accurate species identification within elasmobranchs, which is vital for their conservation and management. Further research addressing these gaps could yield significant improvements in our understanding of elasmobranch biodiversity.

## Conclusion

In conclusion, this research has provided valuable insights into the genetic diversity, distribution, and species relationships of Elasmobranchs in Sri Lanka. Using advanced genomic tools, including the COI and 12S rRNA gene analyses, we have identified key genetic distinctions between species and clarified the taxonomic relationships of several shark and ray species. The findings highlight the importance of utilizing rapid sequencing technologies, such as those offered by Oxford Nanopore Technology (ONT), to enhance export monitoring and field studies. Furthermore, the lack of reference sequences for few species and 12S rRNA gene underscored the need for ongoing data collection and updates to public genetic databases. Overall, this research lays the groundwork for future conservation efforts that focus on improving monitoring of elasmobranch trade domestically and internationally, to inform effective management strategies for threatened Elasmobranch species in Sri Lanka and the wider region.

## References

1. Stevens J. The effects of fishing on sharks, rays, and chimaeras (chondrichthyans), and the implications for marine ecosystems. ICES Journal of Marine Science [Internet]. 2000 Jun 1;57(3):476–94. Available from: 10.1006/jmsc.2000.0724

2. Cortés E. Life History Patterns and correlations in Sharks. Reviews in Fisheries Science [Internet]. 2000 Jan 1;8(4):299–344. Available from: 10.1080/10408340308951115

3. Dulvy NK, Fowler SL, Musick JA, Cavanagh RD, Kyne PM, Harrison LR, et al. Extinction risk and conservation of the world’s sharks and rays. eLife [Internet]. 2014 Jan 21;3. Available from: 10.7554/elife.00590

4. Cariani A, Messinetti S, Ferrari A, Arculeo M, Bonello JJ, Bonnici L, et al. Improving the conservation of Mediterranean chondrichthyans: The ELASMOMED DNA Barcode Reference Library. PLoS ONE [Internet]. 2017 Jan 20;12(1):e0170244. Available from: 10.1371/journal.pone.0170244

5. Field IC, Meekan MG, Buckworth RC, Bradshaw CJA. Chapter 4 Susceptibility of sharks, rays and chimaeras to global extinction. Advances in Marine Biology [Internet]. 2009 Jan 1;275–363. Available from: 10.1016/s0065-2881(09)56004-x

6. Lucifora LO, Balboni L, Scarabotti PA, Alonso FA, Sabadin DE, Solari A, et al. Decline or stability of obligate freshwater elasmobranchs following high fishing pressure. Biological Conservation [Internet]. 2017 May 5;210:293–8. Available from: 10.1016/j.biocon.2017.04.028

7. Dulvy NK, Pacoureau N, Rigby CL, Pollom RA, Jabado RW, Ebert DA, et al. Overfishing drives over one-third of all sharks and rays toward a global extinction crisis. Current Biology [Internet]. 2021 Sep 6;31(21):4773–4787.e8. Available from: 10.1016/j.cub.2021.08.062

8. The global status of sharks, rays, and chimaeras [Internet]. 2024 Jan. Available from: 10.59216/ssg.gsrsrc.2024

9. Convention on International Trade in Endangered Species of Wild Fauna and Flora. Appendices I, II and III. Geneva: CITES Secretariat; 2022 Jun 22. Available from: https://cites.org/sites/default/files/eng/app/2022/E-Appendices-2022-06-22.pdf

10. Peiris MAK, Kumara TP, Ranatunga RRMKP, Liu SYV. Species composition and conservation status of shark from fishery landings and fish markets in Sri Lanka revealed by DNA barcoding. Fisheries Research [Internet]. 2021 Jun 22;242:106045. Available from: 10.1016/j.fishres.2021.106045

11. Blue Resources Trust (BRT). Sri Lanka fisheries data (National). Unpublished data. 2025.

12. Hasarangi DG, Maldeniya R, Haputhantri SS. A Review on shark fishery resources in Sri Lanka. Indian Ocean Tuna Commision–WPEB. 2012:08–15.

13. De Silva ML, Manikarachchi IU. STATUS OF MOBULID GILL PLATES AND SHARK FIN TRADE AT SELECTED SITES OF SRI LANKA. InProceedings of the 2nd Asia International Conference on Multidisciplinary Research 2020 (Vol. 2).

14. Admin. Seizure of dried Sharks fins and fish Gill plates by Biodiversity Protection Branch of Sri Lanka Customs – Sri Lanka Customs [Internet]. Available from: https://www.customs.gov.lk/seizure-of-dried-sharks-fins-and-fish-gill-plates-by-biodiversity-protection-branch-of-sri-lanka-customs.

15. Davidson LNK, Krawchuk MA, Dulvy NK. Why have global shark and ray landings declined: improved management or overfishing? Fish and Fisheries [Internet]. 2015 May 11;17(2):438–58. Available from: 10.1111/faf.12119

16. Hobbs CA, Potts RW, Bjerregaard Walsh M, Usher J, Griffiths AM. Using DNA barcoding to investigate patterns of species utilisation in UK shark products reveals threatened species on sale. Scientific reports. 2019 Jan 31;9(1):1–0.

17. Szyp-Borowska I, Sikora K. DNA barcoding: A practical tool for the taxonomy and species identification of entomofauna. Forest Research Papers [Internet]. 2019 Sep 1;80(3):227–32. Available from: 10.2478/frp-2019-0021

18. Hebert PD, Cywinska A, Ball SL, DeWaard JR. Biological identifications through DNA barcodes. Proceedings of the Royal Society of London. Series B: Biological Sciences. 2003 Feb 7;270(1512):313–21.

19. Vollmer NL, Viricel A, Wilcox L, Moore MK, Rosel PE. The occurrence of mtDNA heteroplasmy in multiple cetacean species. Current Genetics [Internet]. 2011 Jan 13;57(2):115–31. Available from: 10.1007/s00294-010-0331-1

20. Nolan DJ, DaRoza J, Brody R, Ganta K, Luzuriaga K, Huston C, et al. Comparing Gold-Standard Sanger Sequencing with Two Next-Generation Sequencing Platforms of HIV-1 gp160 Single Genome Amplicons. AIDS Research and Human Retroviruses [Internet]. 2024 Jul 3;40(11):659–69. Available from: 10.1089/aid.2024.0012

21. Vasiljevic N, Lim M, Humble E, Seah A, Kratzer A, Morf NV, et al. Developmental validation of Oxford Nanopore Technology MinION sequence data and the NGSpeciesID bioinformatic pipeline for forensic genetic species identification. Forensic Science International Genetics [Internet]. 2021 Mar 12;53:102493. Available from: 10.1016/j.fsigen.2021.102493

22. Parker J, Helmstetter AJ, Devey D, Wilkinson T, Papadopulos AST. Field-based species identification of closely-related plants using real-time nanopore sequencing. Scientific Reports [Internet]. 2017 Aug 10;7(1). Available from: 10.1038/s41598-017-08461-5

23. Pomerantz A, Peñafiel N, Arteaga A, Bustamante L, Pichardo F, Coloma LA, et al. Real-time DNA barcoding in a rainforest using nanopore sequencing: opportunities for rapid biodiversity assessments and local capacity building. GigaScience [Internet]. 2018 Apr 1;7(4). Available from: 10.1093/gigascience/giy033

24. Srivathsan A, Lee L, Katoh K, Hartop E, Kutty SN, Wong J, et al. ONTbarcoder and MinION barcodes aid biodiversity discovery and identification by everyone, for everyone. BMC Biology [Internet]. 2021 Sep 29;19(1). Available from: 10.1186/s12915-021-01141-x

25. Ebert DA, Fowler S, Compagno L. Sharks of the World: A fully Illustrated guide [Internet]. 2013. Available from: http://ci.nii.ac.jp/ncid/BB13693771

26. Last PR, de Carvalho MR, Corrigan SH, Naylor GJ, Séret B, Yang L. The Rays of the World project–an explanation of nomenclatural decisions. Rays of the world. Supplementary information. 2016:1–0.

27. Jabado RW, Ebert DA. Sharks of the Arabian Seas: an identification guide. IFAW, Dubai. 2015.

28. Stevens G, Di Sciara GN, Dando M, Fernando D. Guide to the Manta and Devil Rays of the World.

29. Kemp BM, Monroe C, Smith DG. Repeat silica extraction: a simple technique for the removal of PCR inhibitors from DNA extracts. Journal of Archaeological Science [Internet]. 2006 May 5;33(12):1680–9. Available from: 10.1016/j.jas.2006.02.015

30. Boza JM, Manning JC, Erickson DC. Comparison and optimization of simple DNA extraction methods for LAMP-Based Point-of-Care applications employing submillimeter skin biopsies. ACS Omega [Internet]. 2024 Aug 30;9(37):38855–63. Available from: 10.1021/acsomega.4c05025

31. Drábková L, Kirschner J, Vlĉek Ĉ. Comparison of seven DNA extraction and amplification protocols in historical herbarium specimens of juncaceae. Plant Molecular Biology Reporter [Internet]. 2002 Jun 1;20(2):161–75. Available from: 10.1007/bf02799431

32. Huang W, Xie X, Liang X, Wang X, Chen X. Effects of Different Pretreatments of DNA Extraction from Dried Specimens of Ladybird Beetles (Coleoptera: Coccinellidae). Insects [Internet]. 2019 Mar 29;10(4):91. Available from: 10.3390/insects10040091

33. Abdel-Latif A, Osman G. Comparison of three genomic DNA extraction methods to obtain high DNA quality from maize. Plant Methods [Internet]. 2017 Jan 3;13(1). Available from: 10.1186/s13007-016-0152-4

34. Huanca-Mamani W, Rivera-Cabello D, Maita-Maita J. A simple, fast, and inexpensive CTAB-PVP-Silica based method for genomic DNA isolation from single, small insect larvae and pupae. Genetics and Molecular Research. 2015 Oct;14(3):8001–7.

35. Yue GH, Orban L. Rapid isolation of DNA from fresh and preserved fish scales for polymerase chain reaction. Marine Biotechnology. 2001 May;3:199–204.

36. Ward RD, Zemlak TS, Innes BH, Last PR, Hebert PD. DNA barcoding Australia’s fish species. Philosophical transactions of the royal society B: biological sciences. 2005 Oct 29;360(1462):1847–57.

37. Miya M, Sato Y, Fukunaga T, Sado T, Poulsen JY, Sato K, Minamoto T, Yamamoto S, Yamanaka H, Araki H, Kondoh M. MiFish, a set of universal PCR primers for metabarcoding environmental DNA from fishes: detection of more than 230 subtropical marine species. Royal Society open science. 2015 Jul 1;2(7):150088.

38. Céspedes A, García T, Carrera E, González I, Fernández A, Asensio L, Hernández PE, Martín R. Genetic differentiation between sole (Solea solea) and Greenland halibut (Reinhardtius hippoglossoides) by PCR–RFLP analysis of a 12S rRNA gene fragment. Journal of the Science of Food and Agriculture. 2000 Jan 1;80(1):29–32.

39. Lanfear R, Schalamun M, Kainer D, Wang W, Schwessinger B. MinIONQC: fast and simple quality control for MinION sequencing data. Bioinformatics. 2019 Feb 1;35(3):523–5.

40. Sf A. Basic local alignment search tool. J Mol Biol. 1990;215:403–10.

41. Felsenstein J. Confidence limits on phylogenies: an approach using the bootstrap. evolution. 1985 Jul 1;39(4):783–91.

42. Moron J, Bertrand B, Last PR. A check-list of sharks and rays of western Sri Lanka. Journal of the Marine Biological Association of India. 1998;40(1-2):14257.

43. Fernando D, Bown RM, Tanna A, Gobiraj R, Ralicki H, Jockusch EL, Ebert DA, Jensen K, Caira JN. New insights into the identities of the elasmobranch fauna of Sri Lanka. Zootaxa. 2019 Apr 12;4585(2):201–38.

44. MacNeil MA, Chapman DD, Heupel M, Simpfendorfer CA, Heithaus M, Meekan M, Harvey E, Goetze J, Kiszka J, Bond ME, Currey-Randall LM. Global status and conservation potential of reef sharks. Nature. 2020 Jul 30;583(7818):801–6.

45. Crobe V, Ferrari A, Hanner R, Leslie RW, Steinke D, Tinti F, Cariani A. Molecular taxonomy and diversification of Atlantic skates (Chondrichthyes, Rajiformes): Adding more pieces to the puzzle of their evolutionary history. Life. 2021 Jun 22;11(7):596.

46. Cariani A, Messinetti S, Ferrari A, Arculeo M, Bonello JJ, Bonnici L, Cannas R, Carbonara P, Cau A, Charilaou C, El Ouamari N. Improving the conservation of Mediterranean chondrichthyans: the ELASMOMED DNA barcode reference library. PloS one. 2017 Jan 20;12(1):e0170244.

47. Chang JJ, Ip YC, Neo WL, Mowe MA, Jaafar Z, Huang D. Primed and ready: nanopore metabarcoding can now recover highly accurate consensus barcodes that are generally indel-free. BMC genomics. 2024 Sep 9;25(1):842.

48. Zimmermann J, Hajibabaei M, Blackburn DC, Hanken J, Cantin E, Posfai J, Evans TC. DNA damage in preserved specimens and tissue samples: a molecular assessment. Frontiers in Zoology. 2008 Dec;5:1–3.

49. Boza JM, Manning JC, Erickson DC. Comparison and Optimization of Simple DNA Extraction Methods for LAMP-Based Point-of-Care Applications Employing Submillimeter Skin Biopsies. ACS omega. 2024 Aug 30;9(37):38855–63.

50. Marquina D, Buczek M, Ronquist F, Łukasik P. The effect of ethanol concentration on the morphological and molecular preservation of insects for biodiversity studies. PeerJ. 2021 Feb 12;9:e10799.

51. Nagy ZT. A hands-on overview of tissue preservation methods for molecular genetic analyses. Organisms Diversity & Evolution. 2010 Mar;10:91–105.

52. Collins RA, Trauzzi G, Maltby KM, Gibson TI, Ratcliffe FC, Hallam J, Rainbird S, Maclaine J, Henderson PA, Sims DW, Mariani S. Meta-Fish-Lib: A generalised, dynamic DNA reference library pipeline for metabarcoding of fishes. Journal of Fish Biology. 2021 Oct;99(4):1446–54.

53. Rodríguez MD, Vanhollebeke J, Derycke S. Evaluation of DNA metabarcoding using Oxford Nanopore sequencing for authentication of mixed seafood products. Food Control. 2023 Mar 1;145:109388.

54. Borsa P, Durand JD, Shen KN, Arlyza IS, Solihin DD, Berrebi P. Himantura tutul sp. nov.(Myliobatoidei: Dasyatidae), a new ocellated whipray from the tropical Indo-West Pacific, described from its cytochrome-oxidase I gene sequence. Reports. Biologies. 2013;336(2):82–92.

55. Borsa P. Neotrygon vali, a new species of the blue-spotted maskray complex (Myliobatoidei: Dasyatidae). bioRxiv. 2017 Feb 7:106682.

56. De Bruin GH, Russell BC, Bogusch A. The marine fishery resources of Sri Lanka. FAO;; 1994.

57. Moron J, Bertrand B, Last PR. A check-list of sharks and rays of western Sri Lanka. Journal of the Marine Biological Association of India. 1998;40(1-2):142–57.

58. De Silva RI. Taxonomy and status of the sharks and rays of Sri Lanka. The fauna of Sri Lanka: Status of taxonomy, research and conservation. 2006:294–301.

59. Last PR, White WT. Resurrection of the genus Neotrygon Castelnau (Myliobatoidei: Dasyatidae) with the description of Neotrygon picta sp. nov., a new species from northern Australia. Descriptions of New Australian Chondrichthyans. CSIRO Marine and Atmospheric Research Paper. 2008;22:315–25.

60. Manjaji-Matsumoto BM, Last PR. Two new whiprays, Maculabatis arabica sp. nov. and M. bineeshi sp. nov.(Myliobatiformes: Dasyatidae), from the northern Indian Ocean. Zootaxa. 2016 Jul 28;4144(3):335–53.

61. Lubis K, Sudibyo M, Farajallah A, Hanim N, Renta PP. Identification of Batoid Fishes from North Sumatra waters, Indonesia: Comparing between 12S and 16S rRNA gene as DNA marker. ILMU KELAUTAN: Indonesian Journal of Marine Sciences. 2021 Dec 1;26(4).

