## Supplement documents for "Establishing a DNA reference library for the identification of elasmobranchs in the Indian Ocean using Oxford Nanopore Sequencing"

**Supplements**

**S1 Table.** Description of samples analyzed in this study

| Condition of tissue | Code | Sample details |
| --- | --- | --- |
| Tissues in 95% Ethanol and frozen at -20℃ | E1 | *Mobula mobular* |
|  | E2 | *Carcharhinus falciformis* |
|  | E3 | *Gymnura zonura* |
|  | E4 | *Neotrygon indica* |
|  | E5 | *Himantura uarnak* |
| Salted and dried Tissues at room temperature | D1 | Shark - Piliyandala |
|  | D2 | Shark - Kandy |
|  | D3 | Shark - Gampaha |
|  | D4 | Shark - Valaichchenai |
|  | D5 | Ray - Valaichchenai |
| Frozen fresh tissues | FF1 | *Acroteriobatus variegatus* |
|  | FF2 | *Acroteriobatus variegatus* |
|  | FF3 | *Acroteriobatus variegatus* |
|  | FF4 | *Acroteriobatus variegatus* |
|  | FF5 | Shark jaw tissue |

**S2 Table.** Mean values of DNA concentrations and Purity level with p values obtained from ANOVA test

| **DNA Extraction methods** | **Measures of DNA** | **Sample Types** | | | **p-value** |
| --- | --- | --- | --- | --- | --- |
|  |  | **Tissues in Ethanol (1)** | **Dry Tissues (2)** | **Frozen Tissues (3)** |  |
| **Qiagen (1)** | **Conc 土 SD (ng/ul)** | 141.7土112.3 | 27.4土24.4 | 70.7土26.3 | 0.06 |
|  | **A260/A280 土 SD** | 1.82土0.14 | 1.76土0.23 | 2.05土0.06 | 0.04 |
| **Hot Shot (2)** | **Conc 土 SD (ng/ul)** | 911土1302 | 905土553 | 846土351 | 0.991 |
|  | **A260/A280 土 SD** | 1.58土0.21 | 1.55土0.10 | 1.89土0.05 | 0.005 |
| **chelex (3)** | **Conc 土 SD (ng/ul)** | 155.7土77.3 | 63.4土9.04 | 244.2土82.4 | 0.77 |
|  | **A260/A280 土 SD** | 1.82土0.60 | 1.02土0.20 | 1.47土0.08 | 0.047 |
| **Nucleo Spin (4)** | **Conc 土 SD (ng/ul)** | 138.2土72.6 | 27.2土26.5 | 53.4土48.4 | 0.015 |
|  | **A260/A280 土 SD** | 1.89土0.04 | 1.73土0.38 | 1.87土0.22 | 0.588 |
